# Sodium valproate increases activity of the sirtuin pathway resulting in beneficial effects for spinocerebellar ataxia-3 *in vivo*

**DOI:** 10.1101/2021.03.08.434343

**Authors:** Maxinne Watchon, Luan Luu, Katherine J. Robinson, Kristy C. Yuan, Alana De Luca, Hannah J. Suddull, Madelaine C. Tym, Gilles J. Guillemin, Nicholas J. Cole, Garth A. Nicholson, Roger S. Chung, Albert Lee, Angela S. Laird

## Abstract

Machado-Joseph disease (MJD, also known as spinocerebellar ataxia-3) is a fatal neurodegenerative disease that impairs control and coordination of movement. Here we tested whether treatment with the histone deacetylase inhibitor sodium valproate (SV) prevented a movement phenotype that develops in larvae of a transgenic zebrafish model of the disease. We found that treatment with SV improved the swimming of the MJD zebrafish, increased levels of acetylated histones 3 and 4, but also increased expression of polyglutamine expanded human ataxin-3. Proteomic analysis of protein lysates generated from the treated and untreated MJD zebrafish also predicted that SV treatment had activated the sirtuin longevity signaling pathway and this was confirmed by findings of increased SIRT1 protein levels and sirtuin activity in SV treated MJD zebrafish and HEK293 cells expressing ataxin-3-84Q, respectively. Treatment with resveratrol (another compound known to activate the sirtuin pathway), also improved swimming in the MJD zebrafish. Co-treatment with SV alongside EX527, a SIRT1 activity inhibitor, prevented induction of autophagy by SV and the beneficial effects of SV on the movement in the MJD zebrafish, indicating that they were both dependent on sirtuin activity. These findings provide the first evidence of sodium valproate inducing activation of the sirtuin pathway. Further, they indicate that drugs that target the sirtuin pathway, including sodium valproate and resveratrol, warrant further investigation for the treatment of MJD and related neurodegenerative diseases.

## Background

Machado-Joseph disease (MJD), also known as spinocerebellar ataxia-3, is a fatal neurodegenerative disease characterized by clinical symptoms including ataxia, dystonia, rigidity, muscle atrophy and visual and speech disorder (Costa and Paulson, 2012; Privett and Kardon, 2010; Rub et al., 2013). MJD is the most common of the hereditary ataxias found throughout the world (21-28% of autosomal-dominant ataxia) (Durr et al., 1996; Ranum et al., 1995; Schols et al., 1995), with a high prevalence within the Azores of Portugal (Bettencourt et al., 2008) and Indigenous communities of north east Arnhem Land in Australia (Burt et al., 1996).

MJD is caused by inheritance of an expanded CAG repeat region within the *ATXN3/MJD1* gene on chromosome 14 (Nascimento-Ferreira et al., 2013; Takiyama et al., 1993). In healthy subjects the *ATXN3* gene contains a short CAG trinucleotide repeat region containing 12-40 CAG repeats, but presence of over 40 CAG repeats in this region leads to MJD (Bettencourt et al., 2010; Maciel et al., 1995; Matsumura et al., 1996). This expanded CAG repeat within the *ATXN3* gene results in presence of a long polyglutamine (polyQ) tract towards the C-terminus of the ataxin-3 protein (Maciel et al., 1995; Matsumura et al., 1996).

The ataxin-3 protein is known to function as a deubiquitinating enzyme (DUB) and can also regulate transcription through interactions with other transcriptional coactivators (histone acetyltransferases; HAT) (Li et al., 2002). Studies in models of MJD have demonstrated an interaction between human ataxin-3 (wild-type or mutant) and transcriptional repressor histone deactylase-3 (HDAC; class I HDAC), further highlighting a role for ataxin-3 in regulating transcription (Evert et al., 2006; Feng et al., 2018). Several animal models expressing human ataxin-3 containing an expanded polyQ region have shown hypoacetylation of histones 3 and 4 (Chou et al., 2011; Chou et al., 2014; Lin et al., 2014; Yi et al., 2013) and increased HAT inhibition, suggesting that the expansion of the polyQ tract within the ataxin-3 protein may enhance HDAC activity (Chou et al., 2014; Li et al., 2002).

Whilst there is currently no effective treatment for MJD, a range of recent studies have indicated therapeutic candidates with potential benefit. One candidate that has been trialed on *in vitro, in* vivo and in MJD patients is the class I and IIa HDAC inhibitor sodium valproate (SV, or valproic acid) (Esteves et al., 2015; Lei et al., 2016; Lin et al., 2014; Yi et al., 2013). Similarly, divalproex sodium (comprised of a combination of sodium valproate and valproic acid together) attenuates cytotoxicity in cellular models of MJD (Wang et al., 2019; Wang et al., 2018).

SV has FDA approval for the treatment of epilepsy and bipolar disorder, where it is thought to have a beneficial effect through blocking voltage gated sodium channels and increasing γ-aminobutyric acid (GABA) neurotransmission respectively (Chiu et al., 2013; Macdonald and Kelly, 1995). SV has also been investigated for a range of other neurodegenerative diseases, with clinical trials occurring in Huntington’s disease, Alzheimer’s disease, amyotrophic lateral sclerosis and MJD (Fleisher et al., 2011; Lei et al., 2016; Piepers et al., 2009; Saft et al., 2006). Whilst multiple studies have demonstrated neuroprotective effects of SV treatment, the mode-of-action of this neuroprotective effect is still poorly understood.

To gain a greater understanding of the potential benefits of treatment with SV, we have trialed its treatment on our transgenic zebrafish model of MJD (Watchon et al., 2017). We then carried out label-free proteomics, immunoblot analysis and microscopy to determine the potential mechanisms and cellular pathways by which SV treatment improved movement in the transgenic MJD zebrafish, to identify a pathway for future investigation for the treatment of MJD.

## Results

### Transgenic MJD zebrafish carrying expanded polyQ length exhibit histone hypoacetylation

Immunoblot analysis confirmed that the transgenic zebrafish expressed full-length human ataxin-3 protein of appropriate sizes (72 kDa and 84 kDa for EGFP-Ataxin-3 23Q and 84Q, respectively), as well as endogenous zebrafish ataxin-3 (34 kDa) (Figure 1A). Probing the immunoblots for acetylated histone 3 (H3K9) and 4 (H4K5) revealed a decrease in both with increasing ataxin-3 polyQ length (Figure 1A), with significantly decreased levels of both acetylated H3K9 and acetylated histone 4 (H4K5) in EGFP-Ataxin-3 84Q larvae compared to non-transgenic animals (Figure 1B-C).

**Figure 1.**
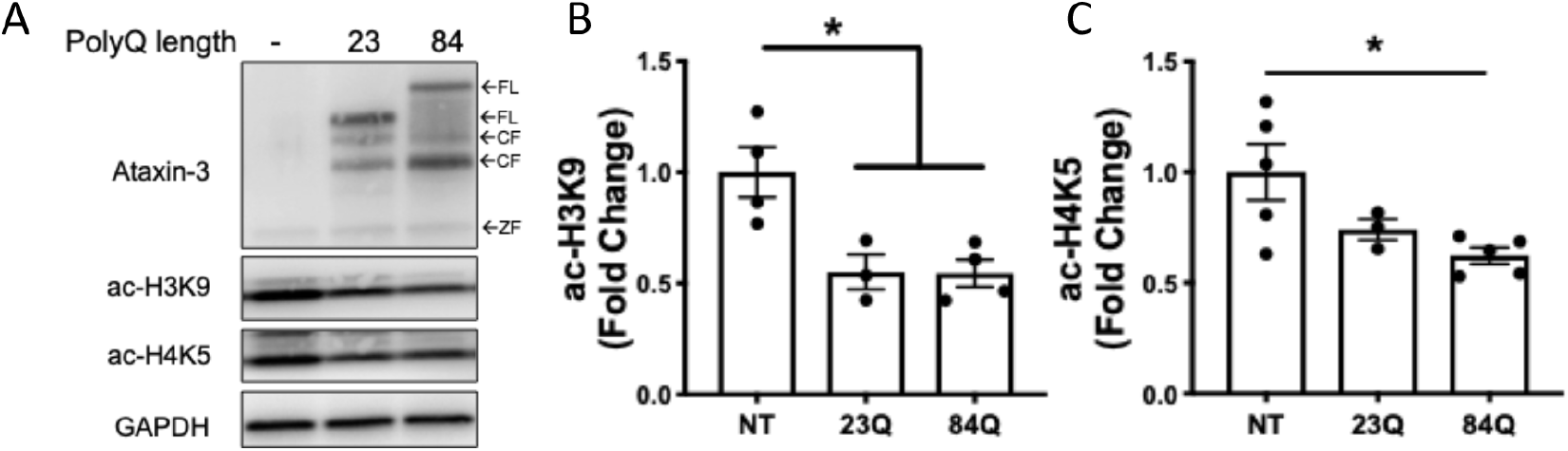
Transgenic MJD zebrafish exhibit hypoacetylation in histones 3 and 4. A) Immunoblots of 6-day-old transgenic MJD zebrafish expressed human ataxin-3 with full-length (FL) ataxin-3, cleaved (CF) ataxin-3 and endogenous zebrafish ataxin-3 (ZF). Transgenic MJD zebrafish also have decreased acetylated histone 3 (H3K9) and 4 (H4K5) expression. B) Quantification of H3K9 showed wild-type and mutant ataxin-3 larvae with significantly lower levels of H3K9 compared to the non-transgenic control (*p<0.022, n=3-5). C) Densitometric analysis of H4K5 expression revealed a significant decrease in expression in the mutant ataxin-3 zebrafish compared to the non-transgenic control (p=0.026, n=3-5). Data is mean ± SEM and statistical analysis used was a one-way ANOVA followed by Tukey post-hoc analysis.

### Sodium valproate improves motor function of mutant ataxin-3 zebrafish and increases levels of acetylated histones

We treated transgenic zebrafish with two doses of the HDAC inhibitor sodium valproate, SV (3.125 μM and 6.25 μM) from 1-6 dpf and then performed motor behaviour testing and whole-body protein extraction for immunoblotting analysis. EGFP-Ataxin-3 84Q larvae swam shorter distances than both non-transgenic and EGFP-Ataxin-3 23Q controls, (Figure 2A), as described previously (Watchon et al., 2017). The impaired swimming of EGFP-Ataxin-3 84Q larvae was rescued by treatment with low-dose SV (3.125 μM), resulting in increased swimming distance. A higher concentration of SV (6.25 μM) did not improve swimming and in fact worsened it, suggesting that this higher drug concentration is reaching toxic levels. Accordingly, we identified an increased rate of morphological abnormalities in animals treated with the higher concentration (data not shown).

**Figure 2.**
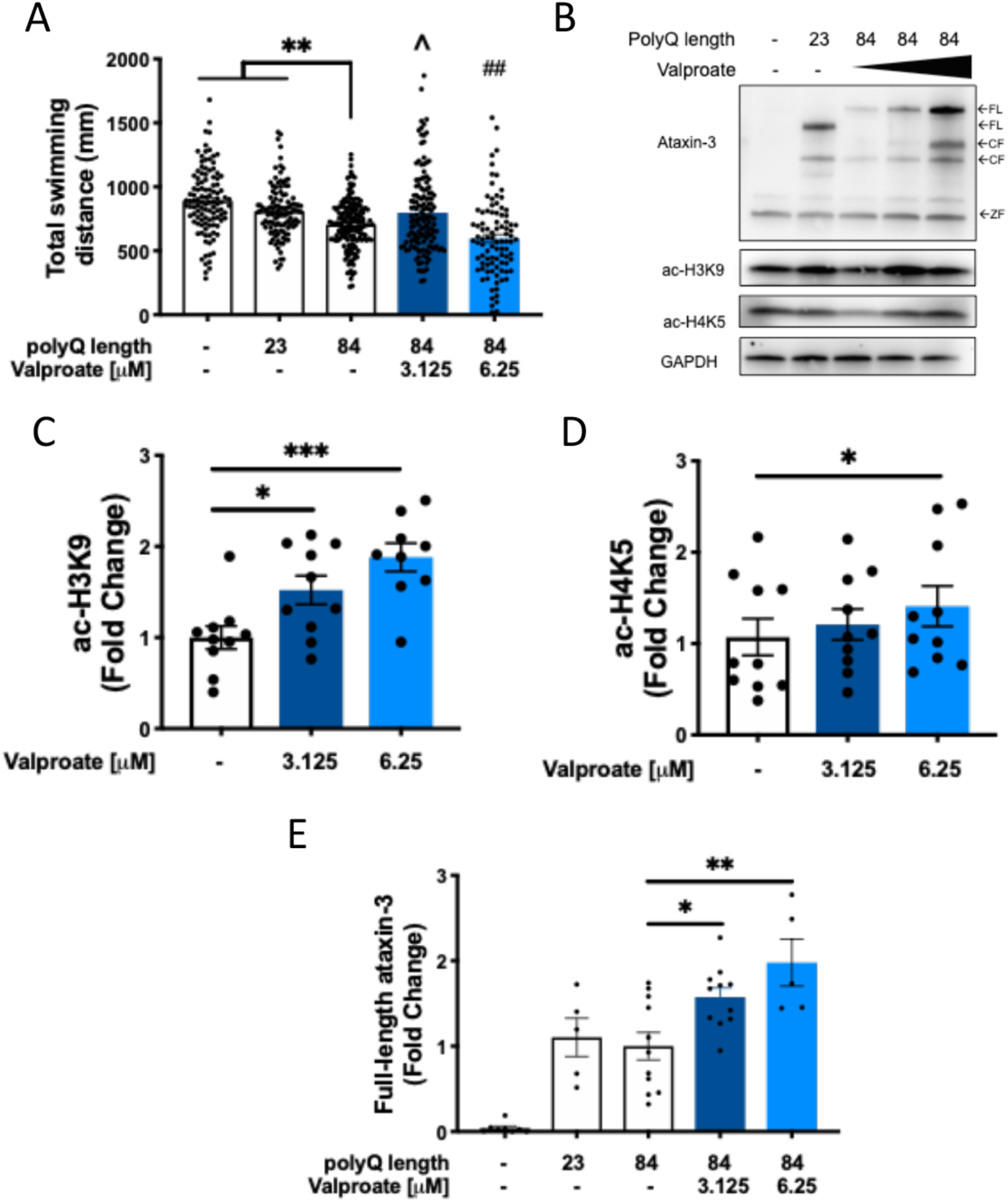
Low-dose sodium valproate treatment improves the swimming of zebrafish larvae expressing polyQ expanded human ataxin-3. A) Treatment of MJD zebrafish with low-dose sodium valproate (SV; 3.125 μM) increased the distance travelled during movement tracking to control treated EGFP-Ataxin-3 84Q (p=0.019, n=93-161). By contrast, high dose SV (6.25 μM) decreased the distance travelled compared to EGFP-Ataxin-3 84Q, ##; p=0.008. B) Western blotting of SV treated EGFP-Ataxin-3 84Q larva resulted in increased levels of full-length human ataxin-3, acetylated histone 4 (at lysine 5, H4K5) and acetylated histone 3 (at lysine 9, H3K9). C) Quantification of H3K9 levels revealed SV treatment increased H3K9 in a dose dependent manner (**p=0.004 and ***p=0.003; n=9). D) Quantification of H4K5 showed an increase with 6.25 μM SV treatment (p=0.043, n=10). E) Quantification of human full-length ataxin-3 (FL) showed significantly increased levels with both concentrations of SV treatment compared to vehicle treated mutant ataxin-3 fish (*p=0.0495, **p=0.002, n=5-11). CF – cleavage fragment, ZF – zebrafish. Data represent mean ± SEM. Statistical analysis used in this figure were paired and unpaired one-way ANOVA with Tukey post-hoc analysis.

Immunoblot analysis of whole-body lysates extracted from the various groups revealed that SV treatment rescued the levels of acetylated histones 3 (H3K9) and 4 (H4K5) (Figure 2B). SV treatment produced a dose dependent increase in acetylated H3K9 levels compared to EGFP-Ataxin-3 84Q receiving vehicle treatment (Figure 2C), with high dose SV causing hyperacetylation of H3K9 compared to non-transgenic and EGFP-Ataxin-3 23Q controls (data not shown). High dose SV also produced an increase in levels of acetylated H4K5 (Figure 2D). We also identified that SV treatment produced a dose dependent increase in the amount of full-length human ataxin-3 present compared to in vehicle treated EGFP-Ataxin-3 84Q zebrafish (up to nearly 2-fold for high dose SV; Figure 2E).

We then examined the effect of SV treatment on HEK293 cells stably expressing human ataxin-3 84Q. Western blotting confirmed the expression of human ataxin-3 84Q (Figure 3A; approximately 52kDa respectively). SV treatment (3 mM) produced significant increases in both acetylated H3K9 and H4K5 levels (Figure 3B-C) and full-length human ataxin-3 levels, compared to vehicle controls (Figure 3D), in a similar manner to the zebrafish findings.

**Figure 3.**
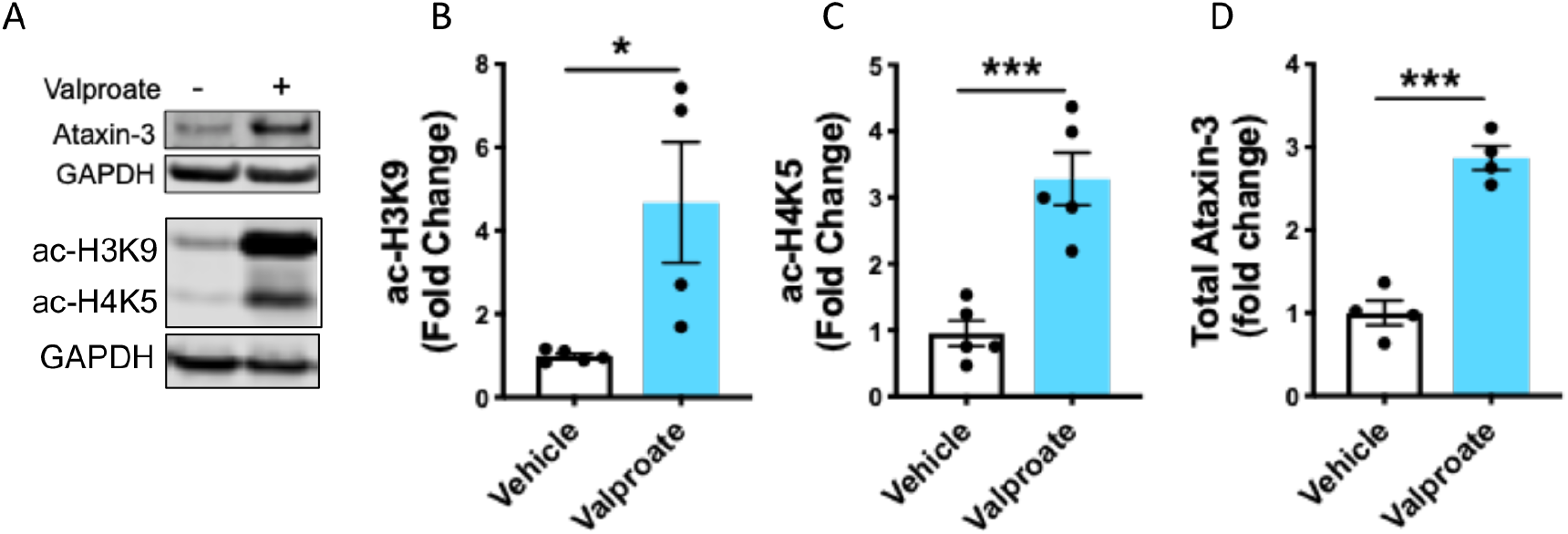
Sodium valproate treatment increases histone acetylation in HEK293 cells expressing human ataxin-3. A) HEK293 cells expressing human ataxin-3 were treated with either sodium valproate (SV) or vehicle and protein lysates extracted from groups of larvae underwent immunoblot analysis for acetylated histones 3 and 4 and human ataxin-3. Quantification revealed that SV treatment increased the amount of B) acetylated histone 3 (H3K9); C) acetylated histone 4 (H4K5); and D) expressed human ataxin-3, compared to vehicle control treatment (p=0.023, p=0.0007, p=0.0001 respectively; n=4-5). Data represents mean ± SEM and statistical analyses used were unpaired student t-tests.

### Unbiased label-free quantitative proteomics identifies various cellular pathways affected by sodium valproate

Our finding of improved swimming following SV treatment, together with increased expression of polyQ-expanded human ataxin-3, was surprising because we hypothesized that increased expression of polyQ-expanded human ataxin-3 would increase the severity of disease rather than improving motor function. This suggested that SV treatment may also induce transcriptional activation of neuroprotective pathways that counteract the negative effects of increased levels of mutant ataxin-3. To identify the downstream molecular pathways that may be responsible for the improved movement of the MJD zebrafish we performed unbiased label-free quantitative proteomics to compare samples from transgenic zebrafish expressing ataxin-3-84Q treated with low-dose SV (3.125 µM) or vehicle control. From triplicate analysis, we identified 1176 and 1209 proteins from vehicle and treated zebrafish respectively, with 984 (70.2 %) proteins identified in common. We therefore identified 192 (13.7%) and 225 (16.1%) unique proteins between the vehicle and SV treated zebrafish lysates (Figure 4A). We carried out gene ontology (GO) annotation of the unique protein lists between the two conditions, and found that with SV treatment, there were less proteins identified that were categorized as carrier proteins, whilst there were more proteins identified that were categorized as having transferase, hydrolase and calcium and nucleic acid binding function (Figure 4B). Overall, there did not appear to be obvious differences in the protein classes that were identified between the vehicle and treated samples.

**Figure 4.**
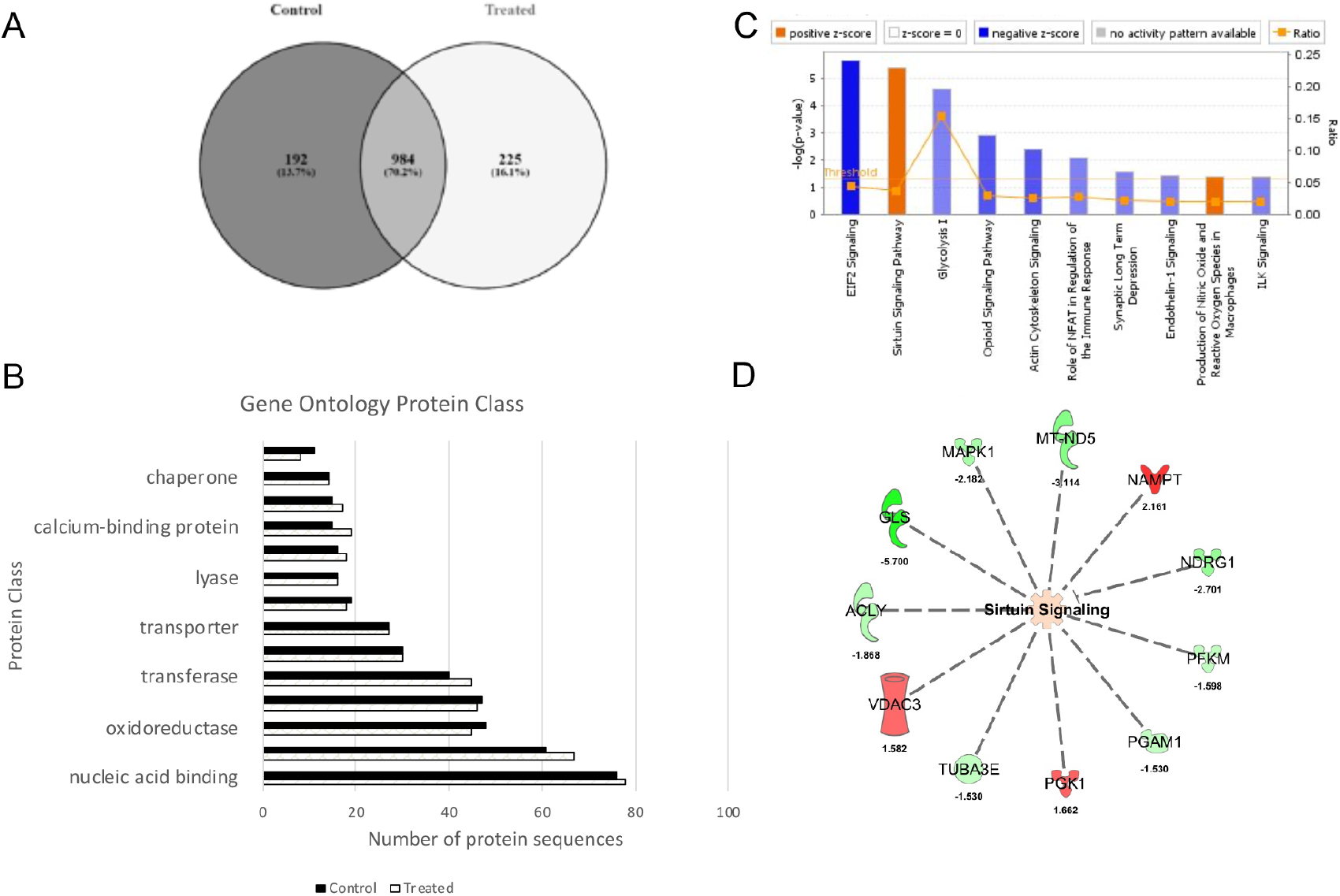
Label-free quantitative proteomics of Ataxin-3 84Q transgenic zebrafish treated with vehicle and sodium valproate. A) Triplicate analyses of vehicle and SV treated transgenic zebrafish identified common and unique proteins. B) Gene ontology (GO) annotation revealed small differences in the identification of proteins, with more proteins (9%) with hydrolase function, while less proteins (3%) were categorized to have carrier activity. C) Ingenuity Pathway Analysis (IPA) predicted activation and inhibition of Sirtuin signaling and EIF2 signaling pathways respectively upon SV treatment of transgenic zebrafish expressing EGFP-Ataxin-3 84Q. Blue indicates IPA predicted inhibition and orange indicates predicted activation of categorised biological function and canonical pathways. D) Predicted upregulation and downregulation of proteins associated with the sirtuin signaling pathway. Green indicates downregulation (0.67-fold) and red indicates upregulation (1.5-fold) of proteins in SV treated EGFP-Ataxin-3 84Q zebrafish compared to the vehicle controls.

To determine differential (quantitative) protein expression between vehicle and SV-treated transgenic zebrafish, we carried out label-free proteomics using normalized spectral abundance factors (NSAF) as described previously (Hogan et al., 2017). We identified approximately 61 downregulated (<0.67-fold) and 86 upregulated (>1.5-fold) proteins in SV-treated zebrafish compared to the vehicle controls (Supplementary 1). Ingenuity Pathway Analysis (IPA) was used to predict activation and/or inhibition of canonical pathways and processes and it predicted the activation of the sirtuin signalling pathway (Z-score = 1, p-value=4.0 × 10^−^6; Figure 4C). Experimental values of proteins associated with the sirtuin pathway (e.g. ACLY, GLS, MAPK1, MT-ND5, NAMPT, NDRG1, PFKM, PGAM1, PGK1, TUBA3E and VDAC3) were obtained from the quantitative analysis (Figure 4D). Based upon the Ingenuity curated database, other biological networks and functions predicted to be affected by SV treatment included the inhibition of “degeneration of neurons” (Z-score = −1.062, p-value = 2.2 × 10^−4^), “apoptosis” (Z-score = −1.696, p-value = 3 × 10^−5^) (Supplementary 2) and inhibition of the EIF2 signaling pathway (Z-score = −1.89, p-value=2.21 × 10^−6^, Figure 4C, Supplementary 1).

### SV treatment induces increased activity of the sirtuin pathway, which has protective effects for MJD zebrafish motor function

LFQ and IPA analysis of canonical signaling pathways of SV treatment on EGFP-ataxin-3-84Q zebrafish predicted an upregulation of the sirtuin pathway. We firstly validated whether SV treatment activated the sirtuin pathway by treating HEK293 cells expressing mutant ataxin-3 (84Q) with SV. Immunoblots of these samples revealed an increase in SIRT1 band intensity compared to the vehicle treatment (Figure 5A-B). As SIRT1 is a deacetylase, we also examined the level of acetylation of p53, a known substrate of SIRT1 deacetylation. Immunoblotting revealed a decrease in acetylated p53 levels upon SV treatment compared to the vehicle control (Figure 5C-D).

**Figure 5.**
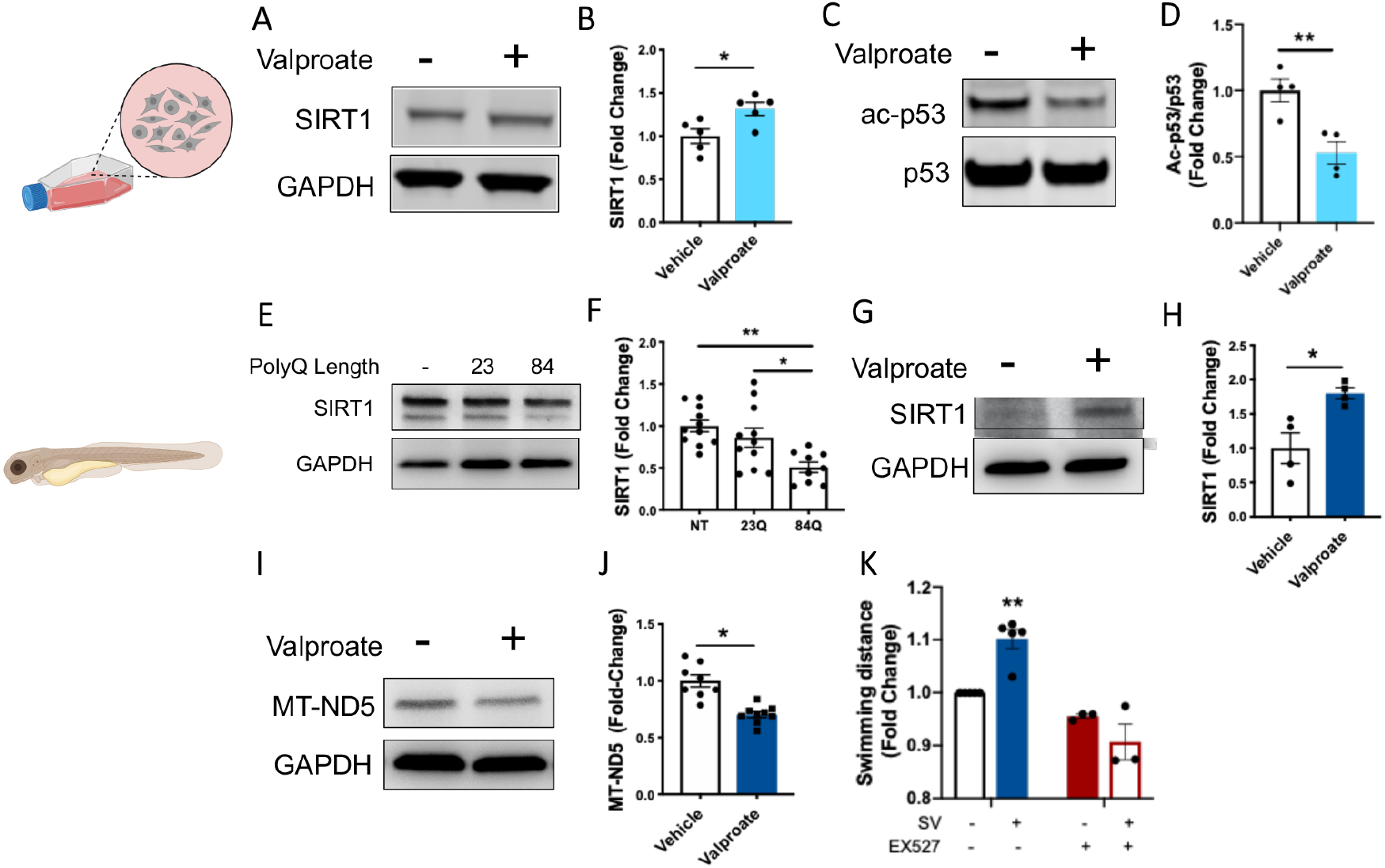
Sodium valproate treatment increases SIRT1 levels and signs of sirtuin activity, validating the increased sirtuin activity predicted by mass spectrometry. A) Immunoblot analysis of ataxin-3 84Q expressing cells treated with SV revealed that SV increases SIRT1. B) Quantification SIRT1 levels revealed a significant increase in SIRT1 from SV treatment (p=0.024, n=5). C) Immunoblot of acetylated p53 and p53, as p53 deacetylation is a marker of sirtuin activity from ataxin-3 84Q expressing HEK293 cells treated with and without SV. D) Quantification of acetylated p53 levels, normalized to p53 levels, revealed that SV treatment resulted in increased p53 deacetylation (p=0.008, n=4). E) Immunoblots of 6-day old transgenic MJD zebrafish show levels of SIRT1. F) Densitometric analysis revealed decreased levels of SIRT1 in zebrafish expressing ataxin-3 with polyQ expansion compared to wild-type ataxin-3 and non-transgenic fish (p=0.025 and p=0.002 respectively, n=9-11). G) Immunoblots of mutant ataxin-3 zebrafish treated with SV showed an increase in SIRT1 expression. H) Quantification of SIRT1 levels revealed an increase with SV treatment (*p=0.015, n=4). I) SV treatment predicted decreased MTND5 levels from the proteomic analysis. Immunoblot of MTND5 showed SV treated EGFP-ataxin-3 84Q protein lysates at 6dpf were decreased. J) Quantification of MTND5 levels confirmed this finding (*p<0.001, n=8-9). K) Whilst treating the EGFP-ataxin-3 84Q zebrafish with SV resulted in the zebrafish swimming longer distances, co-treatment with SV and EX527, or EX527 alone, did not result in increased swimming distances (*p<0.002; SV group significantly greater distances swum than all other groups). Data represents mean ± SEM. Comparisons between vehicle and SV treatment were analysed statistically using unpaired student t-tests and the comparison between ATXN3 genotypes were analysed using an unpaired one-way ANOVA followed by a Tukey post-hoc analysis.

Secondly, we compared the levels of SIRT1 present in the different transgenic ataxin-3 zebrafish genotypes at 6 dpf through immunoblotting. We found there was a decrease in SIRT1 levels in zebrafish expressing ataxin-3 84Q compared to the those expressing wild-type human ataxin-3 and non-transgenic zebrafish (Figure 3E-F). We also validated the proteomic findings by performing immunoblot analysis on lysates from the EGFP-Ataxin-3 84Q zebrafish treated with SV or vehicle control. We found that SIRT1 levels were indeed increased by 1.7-fold in SV treated EGFP-Ataxin-3 84Q samples compared to the vehicle controls (Figure 5G-H). Further suggestion of an increase in activity of the sirtuin pathway was a predicted decrease in the level of mitochondrially encoded NADH Subunit 5 (MT-ND5, or ubiquinone) protein in SV treated samples (Figure 5I-J).

Lastly, we co-treated the mutant ataxin-3 zebrafish with both SV and the SIRT1 inhibitor EX527 (12.5 μM) to determine whether the beneficial effect of SV on zebrafish swimming were indeed through activity of the sirtuin pathway. Treating the MJD zebrafish with SV resulted in an increase in the distance swum by the zebrafish, compared to vehicle treated MJD zebrafish. Co-treatment with SV and EX527 prevented that increase in swimming distance, indicating that the beneficial effect of SV on the MJD zebrafish was dependent on activity of the sirtuin pathway (Figure 5K).

### Confirmation of protective effect of activating sirtuin pathway in MJD zebrafish

To confirm that increased activity of the sirtuin pathway can improve movement of the transgenic MJD zebrafish, we treated our mutant ataxin-3 zebrafish with resveratrol, a natural product known to also activate the sirtuin pathway (Cunha-Santos et al., 2016; Howitz et al., 2003). Treating the mutant ataxin-3 zebrafish with resveratrol (50 μM) for five days significantly increased the distances swum by the zebrafish larvae compared to those treated with vehicle control (Figure 6A). Resveratrol treatment also produced a significant increase in levels of SIRT1 protein compared to those receiving vehicle control (Figure 6B, C). Interestingly, whilst immunoblots of resveratrol treated zebrafish did not show differences between acetylated histone 3 (Figure 5D-E), there was an indication of increased levels of acetylated histone 4 (Figure 5F) and full-length human ataxin-3 (Figure 6G).

**Figure 6.**
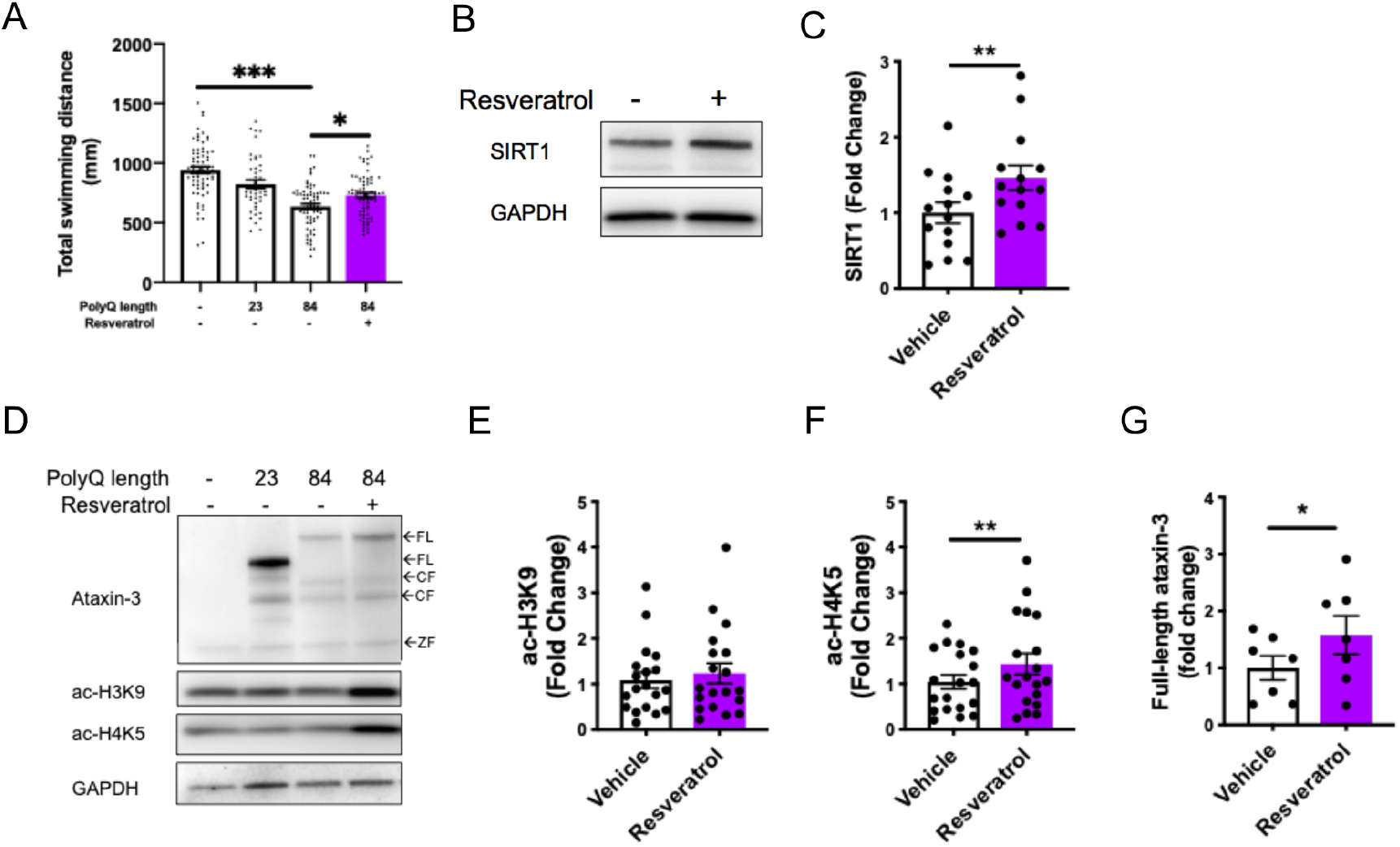
Resveratrol treatment alleviates motor dysfunction whilst simultaneously increasing acetylated histone and SIRT1 levels. A) Mutant ataxin-3 zebrafish at 6 days post fertilization (dpf) showed a decrease in the distance swum (***p<0.001) whilst Resveratrol treatment (50 μM) from 1–6 dpf rescued this dysfunction, (*p=0.0033, n=45-71). B) Immunoblots of vehicle versus resveratrol treated EGFP-Ataxin-3 84Q fish at 6 dpf showed increased SIRT1 levels following resveratrol treatment. C) Quantification revealed that SIRT1 levels were increased by resveratrol (p=0.003, n=14). D) Immunoblots of 6 dpf transgenic MJD zebrafish treated with or without resveratrol increases full-length (FL) human ataxin-3 levels. E) Quantification of levels of acetylated H3K9; F) acetylated H4K5; and G) FL ataxin-3, revealed that resveratrol treatment produced no change in H3K9 levels, an increase in H4K5 (p=0.005) and an increase in FL-ataxin-3 (*p=0.018; n=19, n=19 and n=7 respectively). CF – cleavage fragment, ZF – zebrafish. Data represents mean ± SEM. Statistical analysis used for motor behaviour tracking was a one-way ANOVA followed by a Tukey post-hoc analysis and the immunoblot comparisons were analysed using a paired student t-test.

### Sodium valproate treatment induces macroautophagy in a sirtuin dependent manner

Sodium valproate is a known inducer of activity of the macroautophagy (autophagy) protein quality control pathway (Xia et al., 2016). Increased activity of the sirtuin pathway has also been demonstrated to induce increases in autophagy (Huang et al., 2015; Lee et al., 2008). We therefore examined whether markers of autophagy activity (beclin-1, p62 and LC3II) were elevated following the SV treatment of the MJD zebrafish larvae and HEK293 cells. SV treatment of the MJD zebrafish resulted in increased levels of beclin-1, p62 and LC3II compared to the vehicle-treated control (Figure 7A-D), indicating increased autophagy induction. HEK293 cells stably expressing Ataxin-3 84Q treated with SV had similar levels on p62 to vehicle treated cells (Figure 7E-F) but did show an increase in LC3II levels (Figure 7E, G). We also found that co-treating the MJD zebrafish with SV and EX527 prevented the increased LC3II/I ratio produced by SV treatment alone (Figure 8A-B), suggesting that the increased formation of autophagosomes produced by SV was sirtuin activity dependent. Likewise, co-treating the HEK293 cells expressing Ataxin-3 84Q with SV and EX527 (20 μM) or SV (3 mM) and the autophagy inhibitor 3MA (5 mM), prevented the increased LC3II/I ratio resulting from SV treatment alone (Figure 8C-D), indicating that the induction of autophagy by SV treatment is dependent on SIRT1 activity.

**Figure 7.**
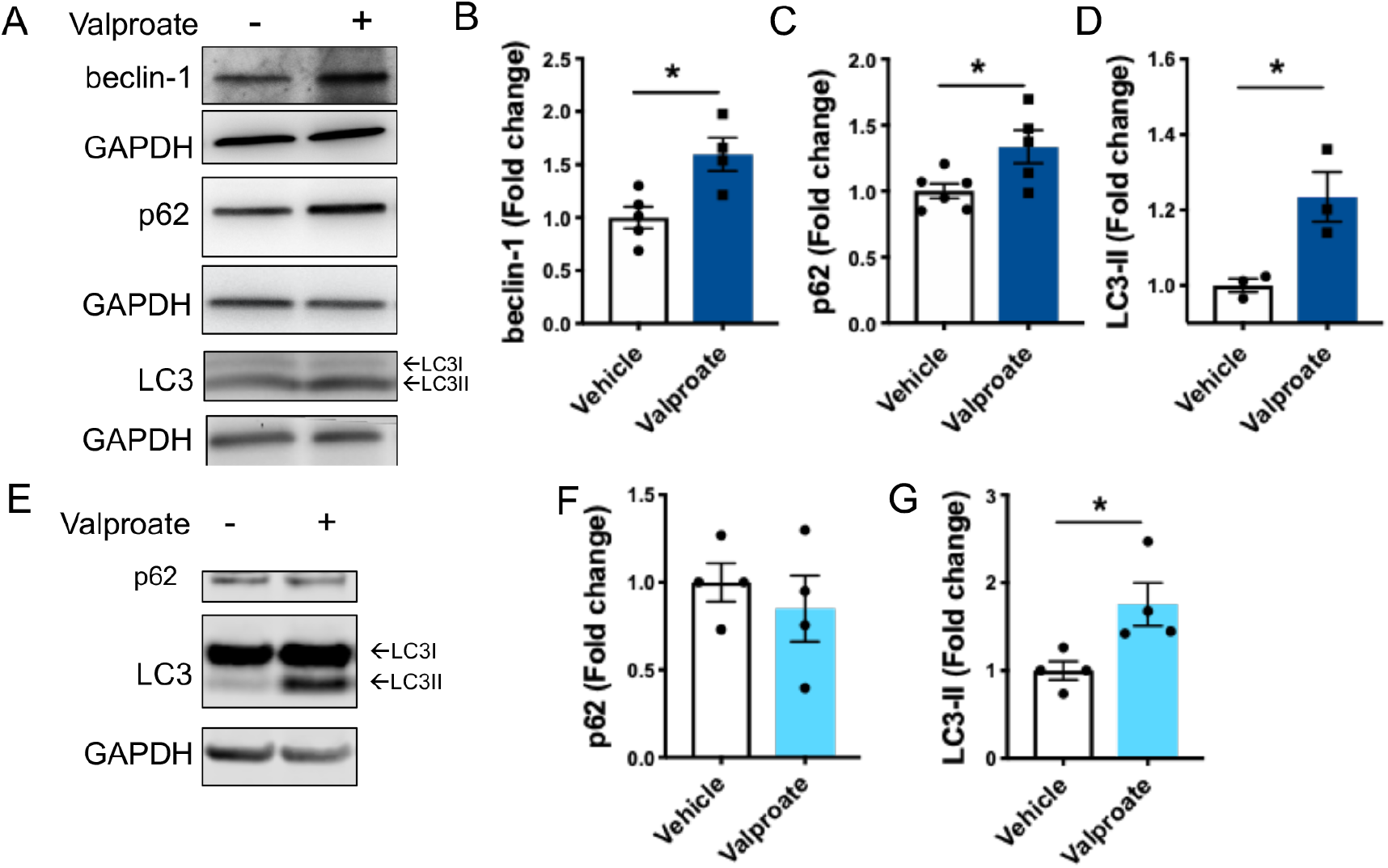
Treatment with sodium valproate (SV) induces activity of the autophagy pathway in transgenic MJD zebrafish and human ataxin-3 expressing HEK293 cells. A) Immunoblots of 6dpf EGFP-Ataxin-3 84Q zebrafish treated with either SV or vehicle control were probed with several markers of the autophagy pathway. Quantification of B) beclin-1, C) p62 and D) LC3-II each revealed a significant increase with SV treatment (*p=0.014, p=0.029 and p=0.026 respectively, n=3-6). E) SV treatment of HEK293 cells expressing Ataxin-3 84Q resulted in similar p62 levels, but increased LC3II levels. Quantification of these substrates revealed F) p62 was not significantly different between SV and vehicle treatment (p=0.518) whilst G) LC3II levels were significantly different for SV compared to vehicle treated (p=0.031, n=4). Comparisons between vehicle and SV treatment were compared using unpaired student t-tests.

**Figure 8.**
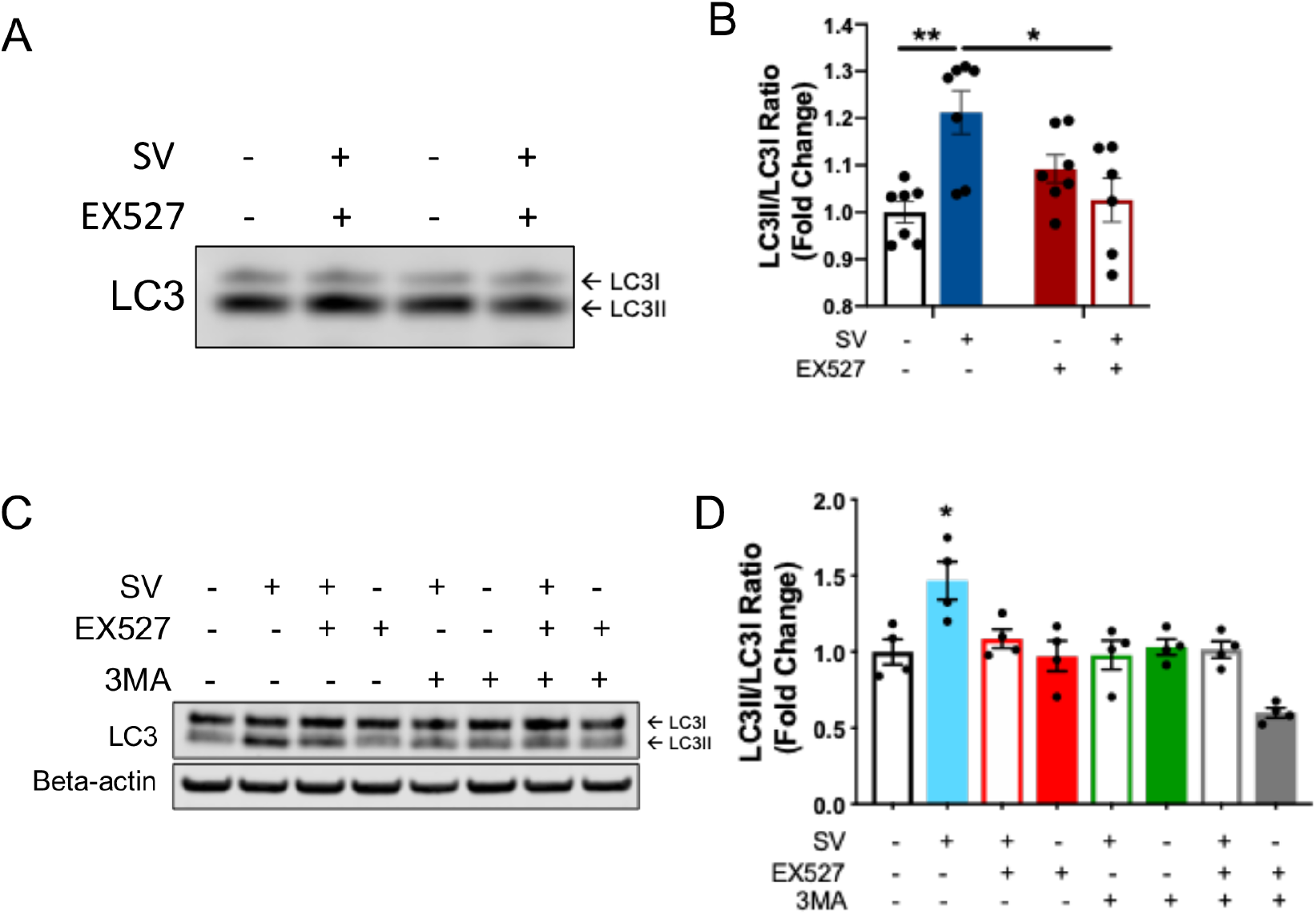
Induction of autophagy following sodium valproate (SV) treatment of MJD zebrafish and human ataxin-3 expressing HEK293 cells is dependent on sirtuin activity. A) Protein lysates from groups of MJD zebrafish larvae underwent immunblotting for LC3B. B) Densiometric analysis revealed that SV treatment increased LC3II/I ratio compared to vehicle treatment (**p=0.003), but co-treatment with SV and EX527 prevented this increase in LC3II/I (*p=0.013, n=6-7). C) Immunoblots for LC3 were performed on lysates from cells treated with SV, EX527 and 3MA, together and alone. I) Densiometric analysis revealed that whilst SV increased LC3II/I ratio, indicating increased autophagosome production, cotreatment with SV and EX527 or SV and 3MA prevented that increase. Data represents mean ± SEM. Comparisons were made using two-way ANOVAs followed by a Tukey post-hoc analysis.

To visualize these changes in LC3 levels, indicative of presence of autophagosomes, we performed immunofluorescent staining of HEK293 cells stably expressing Ataxin-3 84Q treated with SV (3 mM) and/or EX527 (20 μM). It has previously been demonstrated that SIRT1 increases autophagy activity by deacetylating autophagic proteins such as LC3, to allow autophagosomes to relocate from the nucleus to cytoplasm (Huang et al., 2015). Our staining revealed that SV treatment produced a robust increase in the number of LC3 puncta present in the cells, and that EX527 co-treatment prevented that increase (Figure 9A-B). However, whilst the immunostaining images suggested that SV treatment had caused the LC3 to shift to the cytoplasm, manual counting of cells with cytoplasmic LC3 did not detect a statistically significant increase with SV treatment (Figure 9C). Collectively these findings confirm that SV treatment does increase the presence autophagosomes in a SIRT1-dependent manner.

**Figure 9.**
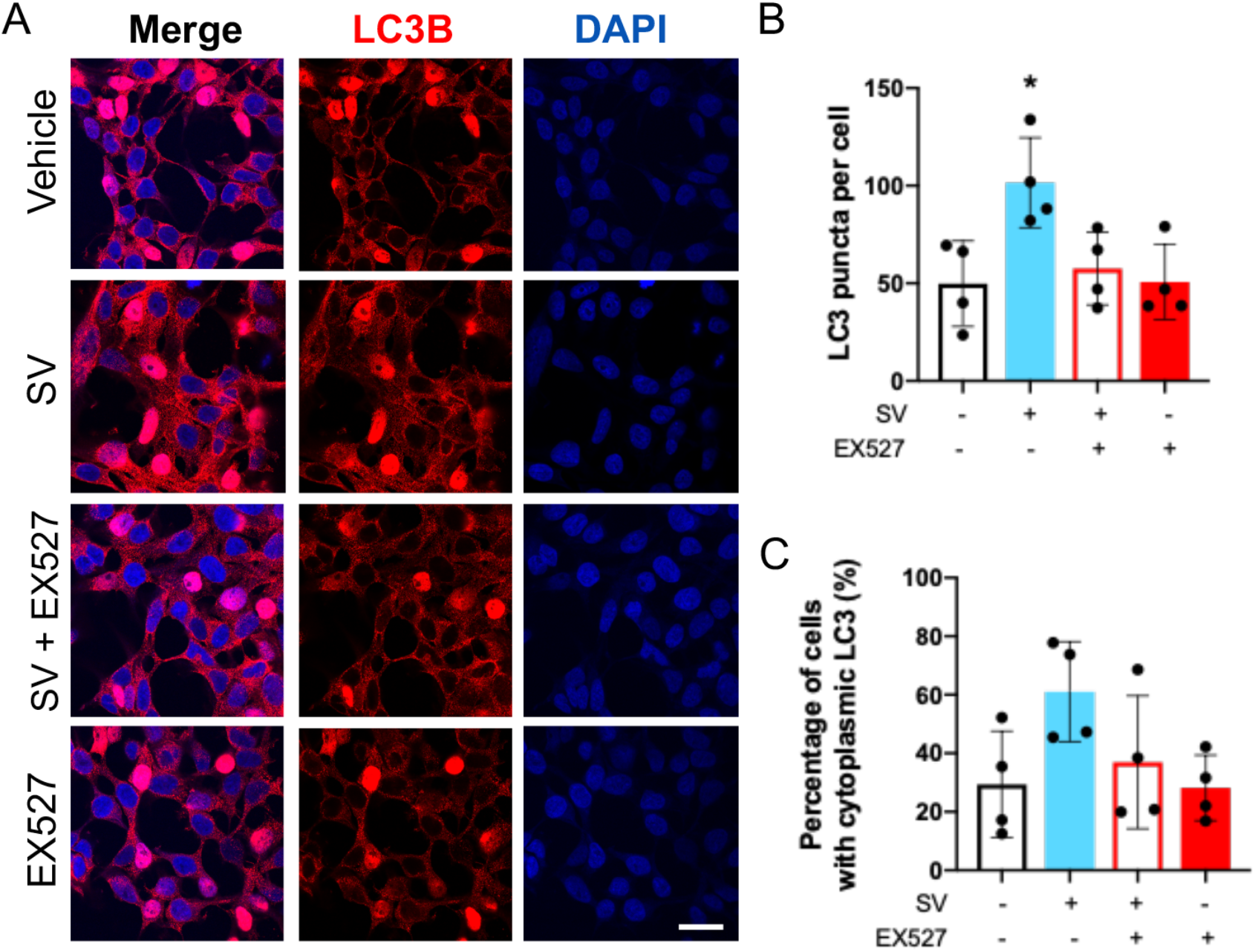
Treatment with sodium valproate (SV) results in an increase in autophagosomes within HEK 293 cells expressing ataxin-3 84Q. A) HEK 293 cells expressing human ataxin-3 84Q were treated with vehicle, sodium valproate (SV), SV and EX527 or EX527 alone, and afterwards stained for LC3 (red) and nuclei (DAPI, blue). Those treated with SV showed strong LC3 (red) staining. B) Automated counting of LC3 (red) puncta revealed that SV treatment increased the presence of LC3 puncta, whilst EX527 cotreatment and EX527 alone had similar numbers of puncta to the vehicle control treated group (*p<0.0483, SV compared to all other groups). C) Manual counting of cells with cytoplasmic LC3 (red) staining revealed that SV had not produced a significant cytoplasmic shift of LC3. Comparisons were analysed using a one-way ANOVA followed by a Tukey post-hoc analysis. Scale bar represents 20 μm.

## Discussion

Here we demonstrate that histone acetylation, which plays a role in transcription regulation, is altered in transgenic MJD zebrafish expressing human ataxin-3 with expanded polyQ tract. Zebrafish expressing EGFP-ataxin-3-84Q exhibited hypoacetylation of histones 3 and 4 compared to non-transgenic zebrafish, suggesting that presence of polyQ expanded ataxin-3 does affect the regulation of histone acetylation by ataxin-3 protein. This finding is in line with previous studies by Yi et al., (2013) and Chou et al., (2011), however it is in contrast with Esteves et al., (2015) and Evert et al., (2006), who found similar or higher levels of histone acetylation, respectively, in cells expressing polyQ expanded ataxin-3.

We used this model, which has benefits of rapid drug testing, to examine whether treatment with sodium valproate, a class I and IIa HDAC inhibitor, would be beneficial for the MJD model. SV has previously been explored as a potential therapeutic agent for MJD (Lei et al., 2016), along with other neurodegenerative diseases such as Huntington’s disease (Zadori et al., 2009), amyotrophic lateral sclerosis (Boll et al., 2014), Parkinson’s disease (Harrison et al., 2015) and Alzheimer’s disease (Qing et al., 2008). Treatment with SV, amongst other HDAC inhibitor compounds, has been demonstrated to positively modify disease phenotypes in other models of MJD including cellular models, *Drosophila* and mice (Chou et al., 2011; Lin et al., 2014; Wang et al., 2018; Yi et al., 2013). In contrast, Esteves et al (2015) have previously reported that valproic acid had limited benefit for a mouse model of MJD. One trial has reported that treating MJD patients with SV is safe and efficacious (Lei et al., 2016). Testing of SV treatment for another polyQ repeat disease, Huntington’s disease, has found a dose dependent alleviation of hyperkinesia, which is hypothesized to be due to an action on GABA transmission (Saft et al., 2006).

In our study, treatment with a low dosage of SV led to improve swimming behavior in zebrafish overexpressing mutant ataxin-3. SV treatment also produced an increase in levels of acetylated histone 3 and high-dose SV increased levels of acetylated histone 4. These results of increased histone acetylation align with previous findings of a beneficial effect of increased histone acetylation following HDAC inhibitor treatment for models of MJD (Lin et al., 2014; Wang et al., 2018; Yi et al., 2013). Importantly, we also found that high concentrations of SV were not protective in our model, and in fact resulted in morphological abnormalities, decreased movement and decreased survival of the zebrafish. This may suggest that there is a dosage threshold for SV treatment and that dose optimisation is an important consideration for future investigations.

Unexpectedly, treatment with SV produced increased expression of human ataxin-3 in the EGFP-ataxin-3 84Q zebrafish and HEK293 cells, perhaps due to an effect of SV on transcription regulation. The ability of SV to provide neuroprotection despite this increase in pathogenic ataxin-3 expression prompted us to explore whether SV additionally activates neuroprotective pathways that alleviate the stresses induced by the expression of mutant ataxin-3. Accordingly, using proteomic protein profiling techniques and pathway analysis, we identified that the SV treatment had produced activation of the sirtuin pathway. This finding was confirmed through immunoblot analysis for SIRT1 in lysates from SV treated zebrafish and HEK293 cells expressing ataxin-3 84Q. We also found that SV treatment of HEK293 cells resulted in decreased levels of acetylated p53, a marker of increased sirtuin activity. We also determined whether SIRT1 protein levels were affected by expression of the polyQ expanded ataxin-3 in the transgenic MJD zebrafish. Indeed, we found that mutant ataxin-3 expression led to decreased levels of SIRT1 compared to the wild-type and non-transgenic animals. This finding is in line with Cunha-Santos and colleagues (2016), wherein mice expressing the mutant form of ataxin-3 contain decreased levels of SIRT1 protein in the cerebellum. Together these findings indicate that treatment with SV can rectify a decrease in SIRT1 levels present in MJD zebrafish.

To test whether this increased sirtuin activity was important for the improvement in the swimming of the MJD zebrafish resulting from SV treatment we performed a co-treatment study involving SV along with the SIRT1 inhibitor, EX527. We found that co-treatment with EX527 prevented the improvement in swimming seen with SV treatment alone, indicating that the protective effect was SIRT1-dependent. Further, we found that treatment with resveratrol, a natural compound known to increase longevity through upregulation of SIRT1 levels, also improved the swimming of the MJD zebrafish. Together these findings indicate that induction of the sirtuin pathway is beneficial for the swimming of the MJD zebrafish.

Our findings of a beneficial effect of SV and resveratrol treatment in our zebrafish model of MJD, along with increasing SIRT1 activity for each, is consistent with Cunha-Santos and colleagues (2016) previous findings that elevating SIRT1 levels in transgenic MJD mice via gene delivery or resveratrol treatment results in decreased neuropathology and motor dysfunction, respectively (Cunha-Santos et al., 2016). A beneficial effect of elevating SIRT1 levels is also in line with previous reports that SIRT1 expression is decreased in MJD models (Cunha-Santos et al., 2016; Wu et al., 2018). This may be explained by previous findings that polyQ-expanded ataxin-3 and huntingtin proteins have been shown to interact with cAMP response element binding (CREB) protein (Toonen et al., 2018), a transcription factor known to regulate levels of SIRT1 (Fusco et al., 2012; Noriega et al., 2011). Interestingly, SV treatment has been shown to activate the CREB pathway (Lin et al., 2014; Sinn et al., 2007), which may explain the increase in SIRT1 levels produced by SV treatment here. Nevertheless, further investigation into the mechanisms by which SV increases activity of the sirtuin pathway is required.

To further explore the mechanisms of the protective effect of SV treatment on the MJD zebrafish, we examined whether markers of autophagy were elevated in the SV treated MJD zebrafish and HEK293 cells. SV treatment has previously been demonstrated to induce activity of the autophagy pathway to produce neuroprotective effects in neurodegenerative disease models (Wang et al., 2015; Zhang et al., 2017). Likewise, increased levels of SIRT1 have been reported to increase autophagic activity, through the removal of the acetyl groups from autophagic substrates Atg5, Atg7 and LC3 (Huang et al., 2015; Lee et al., 2008; Liu et al., 2013). The present study provides evidence of autophagy induction following SV treatment in both the mutant ataxin-3 zebrafish and treated HEK293 cells expressing polyQ expanded ataxin-3. Further, we found that the increased autophagic activity produced by SV treatment in the HEK293 cells and MJD zebrafish were both dependent on SIRT1 activity, as co-treatment of SV and the SIRT1 inhibitor EX527 prevented the induction of autophagy. Analysis of LC3B immunoreactivity in ataxin-3 84Q-expressing HEK293 cells revealed that SV treatment had produced a significant increase in autophagosomes. However, this effect was abolished when SV treatment was combined with EX527, suggesting the increase in autophagosomes, produced by SV treatment was mediated by the sirtuin pathway. Huang et al (2015) had previously reported that SIRT1 overexpression, resveratrol treatment and starvation each cause SIRT1 to deacetylate LC3 driving nuclear export of LC3 to aid formation of autophagosomes. In our study we failed to detect a robust shift of LC3 from the nucleus to the cytoplasm, however this may have been due to imaging limitations.

## Conclusion

In summary, our findings demonstrate for the first time that sodium valproate treatment acts to increase activity of the neuroprotective sirtuin signaling pathway. We identified that an ability of SV to induce increased activity of the autophagy pathway is dependent on activity of the sirtuin activity. Further, our studies identified that SV, along with the sirtuin inducer resveratrol, both improved the movement of our MJD zebrafish. We found that this beneficial effect of SV treatment on the transgenic MJD zebrafish was indeed occurring through activity of the sirtuin pathway. These findings are promising because both sodium valproate and resveratrol have potential to be safe for human treatment as sodium valproate already has FDA approval to treat other neurological conditions and resveratrol is a natural phenol found in the skin of grapes and berries. Further testing of sodium valproate, resveratrol, and other candidates that target the sirtuin pathway, would be beneficial for the development of treatments for MJD, and other related neurodegenerative diseases.

## Methods

### Transgenic zebrafish lines

All animal experiments were performed in accordance with the Animal Ethics Committee of the University of Sydney (K00/2014/575 & 576) and the Animal Ethics Committee of Macquarie University, N.S.W., Australia (ARA: 2016/04 and 2017/019). Zebrafish were housed in a standard recirculating aquarium system at 28.5°C with a 12-hour light and 12-hour dark cycle. Transgenic MJD zebrafish lines were generated as described previously (Watchon et al., 2017), involving crossing of a driver line: Tg(*elavl3*:Gal4-VP16; mCherry) with responder lines Tg(UAS:dsRed,EGFP-ATXN3_Q23) or Tg(UAS:dsRed,EGFP-ATXN3_Q84) to achieve expression of human ataxin-3 containing either 23Q or 84Q. These F1 fish were then in-crossed and their offspring (male and female) were used for the drug treatment studies at the larval stage.

### Motor function testing

All zebrafish behavioral tracking was performed using a ZebraLab Tracking System (Viewpoint) including a ZebraBox for tracking larvae. Tracking of 6-day post fertilization (dpf) larvae was conducted in 24 multi-well plates within a ZebraBox housed with an enhanced light source, under conditions of 6 minutes light, 4 minutes dark and 4 minutes light. The total distance travelled by each larva within the dark phase was calculated.

### Drug treatment studies

Zebrafish embryos (24 hpf) were screened for fluorescence (EGFP and dsRED) indicating that they were positive for the transgenes. Embryos positive for EGFP-Ataxin-3 84Q, of similar expression, were divided into equal numbers and treated for five days with sodium valproate (3.125 μM or 6.25 μM, solubilized in E3 medium), EX527 (12.5 μM, solubilized in DMSO) or resveratrol (50 μM, solubilized in ethanol) through addition of the drug to the E3 buffer that the larvae were incubated in (Nusslein-Volhard and Dahm, 2002). Control treated animals (all genotypes: EGFP-Ataxin-3 23Q, 84Q and non-transgenic control) received the equivalent volume of appropriate vehicle (E3, DMSO or ethanol, respectively). Behavioural testing was performed on morphologically normal larvae at 6 dpf within the ZebraBox.

### Western blotting

Protein lysates were prepared following euthanasia of zebrafish larvae (6 dpf) in RIPA buffer containing protease inhibitors (Complete ULTRA Tablets, Roche), followed by homogenisation via probe sonication. Equal amounts of protein were separated via SDS-PAGE and transferred to PVDF membrane for immunoblot probing. For cell culture lysates, cells were lysed at 3 days in ice-cold RIPA buffer containing protease inhibitors (Roche) and protein concentration was determined by BCA Assay (Gibco BRL, ThermoFisher).

Antibodies used included rabbit anti-MJD (kind gift from H. Paulson), rabbit anti-acetylated histone 3 and 4 (acetylated H3K9 and H4K5 antibodies, Cell Signaling), rabbit anti-MT-ND5 (Abcam), rabbit anti-SIRT1 (Aviva Systems Biology), rabbit anti-beclin-1 (Proteintech), rabbit anti-p62 (MBL Life Science), mouse anti-p62 (Abcam), rabbit anti-LC3B (Abcam), rabbit anti-acetylated p53 (K382; Abcam), rabbit anti-p53 (Abcam), mouse anti-beta-actin (Sigma) and mouse anti-GAPDH (Proteintech). The immunoblots were probed with appropriate secondary antibodies (Promega) and visualised by chemiluminescence (Supersignal detection kit, Pierce) using a BioRad GelDoc System or via fluorescence using the LiCor Odyssey System. The intensities of the bands were quantified by Image Studio Lite and the target protein expression level was determined by normalizing against the loading control protein.

### In-gel trypsin digestion

Whole lysates of 6 dpf EGFP-ataxin-3 84Q (n = 3 for vehicle and SV treatment) were separated by SDS-PAGE for 5 minutes (~1 cm into the gel) and stained with Coomassie Brilliant Blue. Stained proteins were excised from the gel and prepared for in-gel trypsin digestion as described (Shevchenko et al., 2006). Briefly, gel pieces were equilibrated and dehydrated with 50 mM NH_4_HCO_3_ pH 7.8 and 50 mM NH_4_HCO_3_/50% (v/v) acetonitrile pH 7.8 respectively and dried under vacuum centrifugation. The protein gel pieces were reduced with 10 mM DTT at 55°C for 30 minutes and alkylated with 20 mM iodoacetamide (IAA) for 1 hour in the dark at room temperature. The proteins were digested with trypsin (12.5 ng/μl) overnight at 37°C. Following digestion, tryptic peptides were passively diffused in the presence of 50% (v/v) acetonitrile/2 % (v/v) formic acid (2x) in a bath sonicator and collected. The acetonitrile was evaporated by vacuum centrifugation, and tryptic peptides were desalted on a pre-equilibrated C_18_ Sep-Pak cartridge and eluted in 50 % (v/v) ACN, 0.1 % (v/v) formic acid, and dried under vacuum centrifugation.

### Reverse phase (C_18_) liquid chromatography mass spectrometry (LC-MS/MS)

Peptide fractions were separated on a nanoLC system (Thermo) employing a 100-minute gradient (2%–50% v/v acetonitrile, 0.1% v/v formic acid for 95 minutes followed by 90% v/v acetonitrile, 0.1% v/v formic acid for 5 minutes) with a flow rate of 300 nL/min. The peptides were eluted and ionized into Q-Exactive mass spectrometer (Thermo). The electrospray source was fitted with an emitter tip 10μm (New Objective, Woburn, MA) and maintained at 1.8 kV electrospray voltage. FT-MS analysis on the Q-Exactive was carried out with a 70,000 resolution and an AGC target of 1×10^6^ ions in full MS (scan range m/z 350-1800); and MS/MS scans were carried out at 17,500 resolution with an AGC target of 2×10^5^ ions. Maximum injection times were set to 60 and 100 milliseconds respectively. The ion selection threshold for triggering MS/MS fragmentation was set to 15,000 counts and an isolation width of 2.0 Da was used to perform HCD fragmentation with normalised collision energy of 30 %.

Spectra files were processed using the Proteome Discoverer 1.4 software (Thermo) incorporating the Mascot search algorithm (Matrix Sciences, UK) and the *Danio rerio* NCBI RefSeq protein database (05/05/2017, Sequences 46751). Peptide identifications were determined using a 20-ppm precursor ion tolerance and a 0.1-Da MS/MS fragment ion tolerance for FT-MS and HCD fragmentation. Carbamidomethylation modification of cysteines was considered a static modification while oxidation of methionine and acetyl modification on N-terminal residues were set as variable modifications allowing for maximum three missed cleavages. The data was processed through Percolator for estimation of false discovery rates. Protein identifications were validated employing a q-value of 0.01 (1% false discovery rate).

Label-free quantitative proteomics was carried out by calculating the normalised spectral abundance factors (NSAF) according to (Zybailov et al., 2006), which takes into account the length of a given protein as well as the total amount of protein in a given sample. A fraction (0.5) of a spectral count was added to all samples to account for missing values and total spectral counts for at least one condition were set to a minimum of 5. NSAF values were log_2_-transformed and Student’s t-tests were used to identify significant (P ≤0.05) changes in protein abundance between transgenic zebrafish expressing EGFP-ataxin-3 84Q treated with vehicle and SV. Statistical data preparation and tests were done using Microsoft Excel. Enriched GO annotations and signaling pathways were identified using the Database for Annotation, Visualization and Integrated Discovery (DAVID) and Ingenuity^®^ Pathway Analysis (IPA; QIAGEN) to predict activation and inhibition of cellular pathways upon treatment.

The mass spectrometry proteomics data has been deposited to the ProteomeXchange Consortium via the PRIDE (Vizcaino et al., 2016) partner repository with the dataset identifier PXD009612 (Username; Password: ta6siSAA).

### Sodium valproate treatment of HEK293 cells expressing human ataxin-3

Wildtype HEK293 cells were grown in DMEM supplemented with 10% fetal bovine serum and maintained at 37°C and 5% CO_2_. Wild type HEK293 cells were transfected with Lipofectamine LTX with plus reagent (Thermoscientific) and 5 μg of DNA from pcDNA3.1 vectors containing full length human ataxin-3 (containing a polyQ stretch of 28 glutamines or 84 polyglutamines) and a neomycin resistance gene. Cells were treated with 500 μg/mL of Geneticin (Sigma Aldrich), a selective antibiotic analogue of neomycin, to select cells stably expressing pcDNA3.1 vectors. Stable transgenic expression of ataxin-3 or an empty vector control was maintained via treatment with 250 μg/mL Geneticin.

For SV treatments, cells were seeded into 24 well plates at a density of 30,000 cells/cm^2^ and left overnight in a 37 °C incubator supplemented with 5% CO_2_. After 24 hours, cells were treated with SV (3 mM), EX527 (20 μM) or vehicle control in complete growth media. Growth media containing the drug compound was replenished daily for 3 days.

### Immunostaining and analysis of subcellular localization of LC3 in HEK 293 cells expressing human ataxin-3

For immunostaining, coverslips (n = 3 per condition) were fixed with 4% paraformaldehyde (20 minutes) and permeabilised with 0.2% Triton X-100 in PBS. Coverslips were briefly washed (2 × 5 minutes in PBS) and incubated in 2% BSA in PBS to block against non-specific binding and incubated in 2% BSA containing rabbit anti-LC3B primary antibody (Abcam). Coverslips were then washed 3 × 5 minutes in PBS, followed by incubation with goat anti-rabbit Alexa 555 secondary antibody (Life Technologies). Coverslips were washed 3 × 5 minutes in PBS and finally mounted onto glass slides using Prolong Gold mounting medium containing 4′,6-diamidino-2-phenylindole (DAPI) (Thermofisher). Images were acquired via confocal microscopy using a Zeiss LSM-880 confocal microscope (Plan-Apochromat 40x/1.3 Oil DIC UV-IR M27objective, master gain: 800) running Zen Black software (Zeiss, Gottingen, Germany). To visualize LC3 staining, a DPSS561 laser was used (5.6 % laser power) and a UV laser (3% laser power) was used the visualize DAPI-positive nuclei. Images were processed using Airyscan mode and brightness and contrasted was adjusted in an identical manner.

The number of LC3 stained puncta was quantified in an automated manner within ImageJ, through use of the particle analysis function to count LC3 stained (red) particles, with the number of puncta per image divided by the number of DAPI stained (blue) nuclei per image. Manual counting of the number of cells with cytoplasmic LC3-staining, per image, was performed by an experimenter blind to experimental group. The percentage of cells with cytoplasmic LC3-staining was calculated by dividing those counts by the total number of DAPI stained nuclei per image.

### Statistics

Data analysis was performed using GraphPad Prism (version 8). Densitometric analysis of vehicle treatment versus SV (or resveratrol) were compared using a Student t-test. Specific analyses are described in more detail in the figure legend. Remaining analysis involved the use of a one-way ANOVA, followed by a Tukey post-hoc test to identify differences unless otherwise stated. Statistically significant differences are defined as *p<0.05.

## Declarations

### Ethics approval and consent to participate

All animal experiments were performed in accordance with the Animal Ethics Committee of the University of Sydney (K00/2014/575 & 576) and the Animal Ethics Committee of Macquarie University, N.S.W., Australia (ARA: 2016/04).

### Consent for publication

Not applicable.

### Availability of data and material

The datasets produced during the current study are available from the corresponding author on reasonable request.

### Competing interests

The authors declare that they have no competing interests.

### Funding

This work was funded by National Health and Medical Research Council (APP1069235 and APP1146750), the MJD Foundation of Australia and Macquarie University Research Development Grant. The Swedish SCA Network also provided donation support for this work. NJC was supported by The Snow Foundation, MND Association Australia and BitFury.

### Authors’ contributions

MW performed the zebrafish studies, data analysis, interpretation and manuscript preparation. AL, ADL and HJS performed the proteomics and pathway analysis. LL and KJR performed the cell culture experiments and data analysis. KCY and MCT assisted with the zebrafish studies. GJG, NJC, GAN, RC assisted with experimental design and interpretation. ASL designed the experiments, performed data analysis, interpretation and manuscript preparation. All authors read and approved the final manuscript.

## Acknowledgements

The graphical abstract and Figure 5 were prepared using icons from BioRender. The authors would like to thank Henry Paulson for the MJD antibody. The authors also thank the Australian Proteomics Analysis Facility (APAF) established under the Australian Government’s NCRIS program for the mass spectrometry, and staff members (past and present) of the zebrafish facilities at the Brain and Mind Centre and Macquarie University for the care of the zebrafish described within this manuscript.

---

**Supplementary 1.**
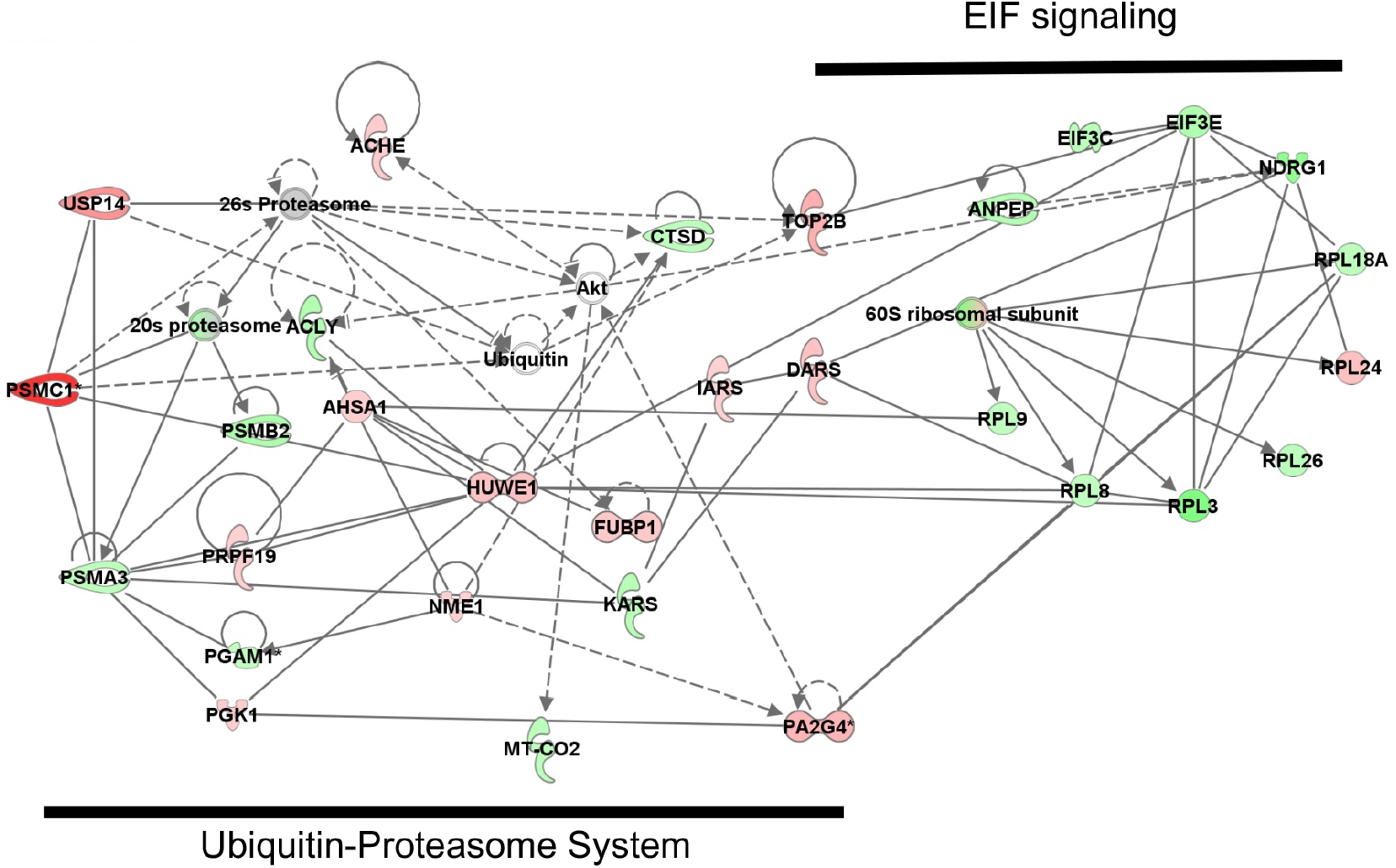
Using label-free quantitative proteomics results, IPA demonstrated clustering of components of EIF signalling and the ubiquitin-proteome system and predicted the inactivation of the EIF2 signalling pathway. Green indicates downregulation (0.67-fold) and red indicates upregulation (1.5-fold) of proteins in SV treated EGFP-Ataxin-3 84Q zebrafish compared to the vehicle controls.

**Supplementary 2.**
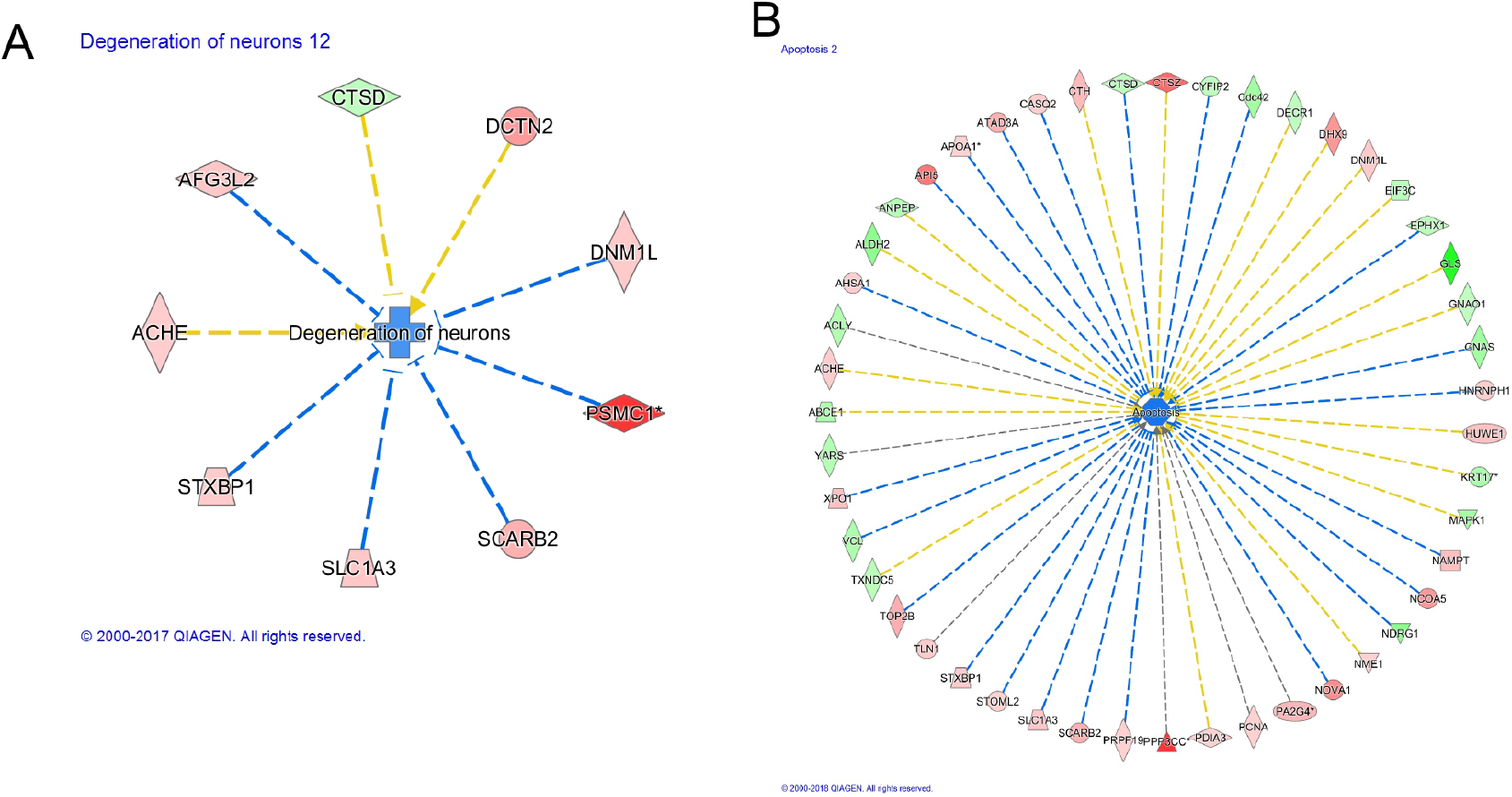
IPA predicted inhibition of biological and disease functions A) “Degeneration of neurons” (Z-score −1.062 p-value= 3.3 × 10–4) and B) “Apoptosis” (Z-score −1.246 p-value=2.03 × 10–5) suggesting cell death processes were suppressed upon valproate treatment. Green indicates downregulation (0.67-fold) and red indicates upregulation (1.5-fold) of proteins in SV treated EGFP-Ataxin-3 84Q zebrafish compared to the vehicle controls. Blue indicates predicted inhibition and orange indicates predicted activation of categorised biological function/pathway.

## Notes

### Competing Interest Statement

The authors have declared no competing interest.

